# Resistance to age-related hearing loss in the echolocating big brown bat (*Eptesicus fuscus*)

**DOI:** 10.1101/2024.07.15.603592

**Authors:** Grace Capshaw, Clarice A. Diebold, Danielle M. Adams, Jack Rayner, Gerald S. Wilkinson, Cynthia F. Moss, Amanda M. Lauer

## Abstract

Hearing mediates many behaviors critical for survival in echolocating bats, including foraging and navigation. Most mammals are susceptible to progressive age-related hearing loss; however, the evolution of biosonar, which requires the ability to hear low-intensity echoes from outgoing sonar signals, may have selected against the development of hearing deficits in echolocating bats. Although many echolocating bats exhibit exceptional longevity and rely on acoustic behaviors for survival to old age, relatively little is known about the aging bat auditory system. In this study, we used DNA methylation to estimate the ages of wild-caught big brown bats (*Eptesicus fuscus*) and measured hearing sensitivity in young and aging bats using auditory brainstem responses (ABRs) and distortion product otoacoustic emissions (DPOAEs). We found no evidence for hearing deficits in aging bats, demonstrated by comparable thresholds and similar ABR wave and DPOAE amplitudes across age groups. We additionally found no significant histological evidence for cochlear aging, with similar hair cell counts, afferent, and efferent innervation patterns in young and aging bats. Here we demonstrate that big brown bats show minimal evidence for age-related loss of peripheral hearing sensitivity and therefore represent informative models for investigating mechanisms that may preserve hearing function over a long lifetime.

## Introduction

Hearing is essential for echolocating bats that rely extensively on their auditory systems to forage, navigate, and avoid obstacles. The evolution of echolocation in bats has been correlated with adaptations at all levels of auditory processing to enable active acoustic sensing of complex and dynamic environments. These include cochlear specializations to enhance detection of behaviorally relevant frequencies (e.g., acoustic foveae in bats that use constant frequency echolocation calls, [1–3] and central processing specializations that facilitate the extraction of fine spectro-temporal cues from received echoes (reviewed in [4]). To receive detectable echo returns from sonar objects, bats emit intense sonar signals that can reach levels as high as 110-140 dB [5–8]. Consequently, echolocating species are exposed to intense self-generated sounds. Further, many species form high-density aggregations, where sonar sounds emitted by other individuals may be potentially damaging to the cochlea. The critical role of hearing in the fitness and survival of echolocating bats suggests that the evolution of this active sensing system may have introduced selective pressures to protect the auditory system from damage over a lifetime of exposure to sound.

The aging auditory system in most mammals shows a progressive loss of hearing sensitivity that begins with high-frequency deficits and extends to low frequencies over time [9–11]. Although the etiology of age-related hearing loss (ARHL) is highly variable, depending on genetic, epigenetic, and environmental factors, its onset is generally correlated with senescent changes to the peripheral structures of the auditory system, including loss of inner and outer hair cells, loss of ribbon synapses and retraction of auditory nerve fibers (i.e., cochlear synaptopathy), and deterioration of the stria vascularis [9,12–18]. The molecular mechanisms underlying ARHL are hypothesized to result from inter-related metabolic and physiological changes over the lifespan that lead to the accumulation of reactive oxygen species and increase susceptibility to cellular dysfunction [19–22]. Recently, bats have emerged as a powerful model for aging studies, due to their extended lifespans relative to comparably sized mammals [23–26]. Extreme longevity in bats is correlated with effective immune responses and reduced susceptibility to oxidative stress [27–32], leading to ‘healthy’ aging characterized by reduced mitochondrial dysfunction and resistance to cellular senescence [33–35]. Bats, therefore, represent a useful model for understanding mechanisms that support cochlear health over a long lifespan.

Echolocation-based foraging and navigation require the ability to hear quiet returning echoes within complex acoustic backgrounds. For bats that survive to old age, hearing deficits could lead to a multitude of negative outcomes, ranging from reduced prey capture success to the inability to detect and avoid obstacles in flight. We hypothesize that echolocation imposes selective pressures to preserve hearing function across the lifespan, especially in species that require echolocation-based active sensing for prey capture. Although bats are not immune to hearing loss [36], and indeed, some species appear vulnerable to ARHL [37], recent evidence indicates that species differences in echolocation behaviors may correlate with differential susceptibility to hearing loss. For example, echolocating bat species have shown evidence for resistance to noise-induced cochlear hair cell damage whereas non-echolocating visually dominant species were susceptible to acoustic overexposure and showed levels of hair cell loss comparable to that observed in mice [38]. Therefore, bats may vary in their susceptibility to cochlear aging in a manner that correlates with their reliance on echolocation cues for survival. To date, ARHL has only been reported in the Egyptian fruit bat [37], a lingual echolocator that shows preference for visual over acoustic cues during behavioral tasks [39] and may not require acute auditory sensitivity across the lifespan compared to insectivorous echolocators that use their hearing to hunt prey.

In this study, we evaluated hearing sensitivity, outer hair cell function, and cochlear morphology in young and aging big brown bats (*Eptesicus fuscus*). *E. fuscus* is an insectivorous laryngeal echolocator that produces high-intensity calls (up to 138 dB SPL at 0.1 m [40]) to pursue and intercept prey in flight. This species has been reported to live up to 19 years in the wild [41,42] (mean longevity: 6 years [43]) and presumably good auditory sensitivity is essential for survival into old age, as foraging efficiency depends critically on the detection of weak, high frequency echoes. Further, *E. fuscus* appears to be resistant to noise-induced hearing damage based on physiological and behavioral testing [44–46]. The specializations that confer resistance to noise damage in this species may enhance survival of cochlear hair cells and their afferent and efferent neurons into old age, which should preserve hearing sensitivity across the lifespan. Here we predicted that aging *E. fuscus* will have peripheral auditory sensitivity comparable to young individuals to facilitate effective echolocation-based behaviors throughout their natural lifespan.

## Results

### Physiological hearing sensitivity does not differ across young and aging bats

We measured hearing sensitivity thresholds by recording auditory brainstem responses (ABRs) in 13 young bats ranging in age from 1.2 to 5.9 years old (mean age: 3.5 years; mean weight: 17.2 g) and 10 aging bats from 6.8 to 12.5 years old (mean age: 8.2 years; mean weight: 17.0 g). *E. fuscus* are sensitive to sounds from 4-84 kHz, with peaks in sensitivity from 12-24 kHz and 60-72 kHz (Figure 1A). ABR-derived audiograms for young and aging *E. fuscus* were similar across the range of frequencies tested, indicating comparable hearing sensitivity across age groups (Figure 1). Auditory detection thresholds were not statistically different across age group (F_1,18.55_= 0.66, p=0.43) or sex (F_1,17.57_= 4.09, p=0.06), nor was there a significant interaction effect of age group with frequency (F_15,181.56_= 0.58, p=0.89) or age group with sex (F_1,17.57_= 1.20, p=0.29).

**Figure 1.**
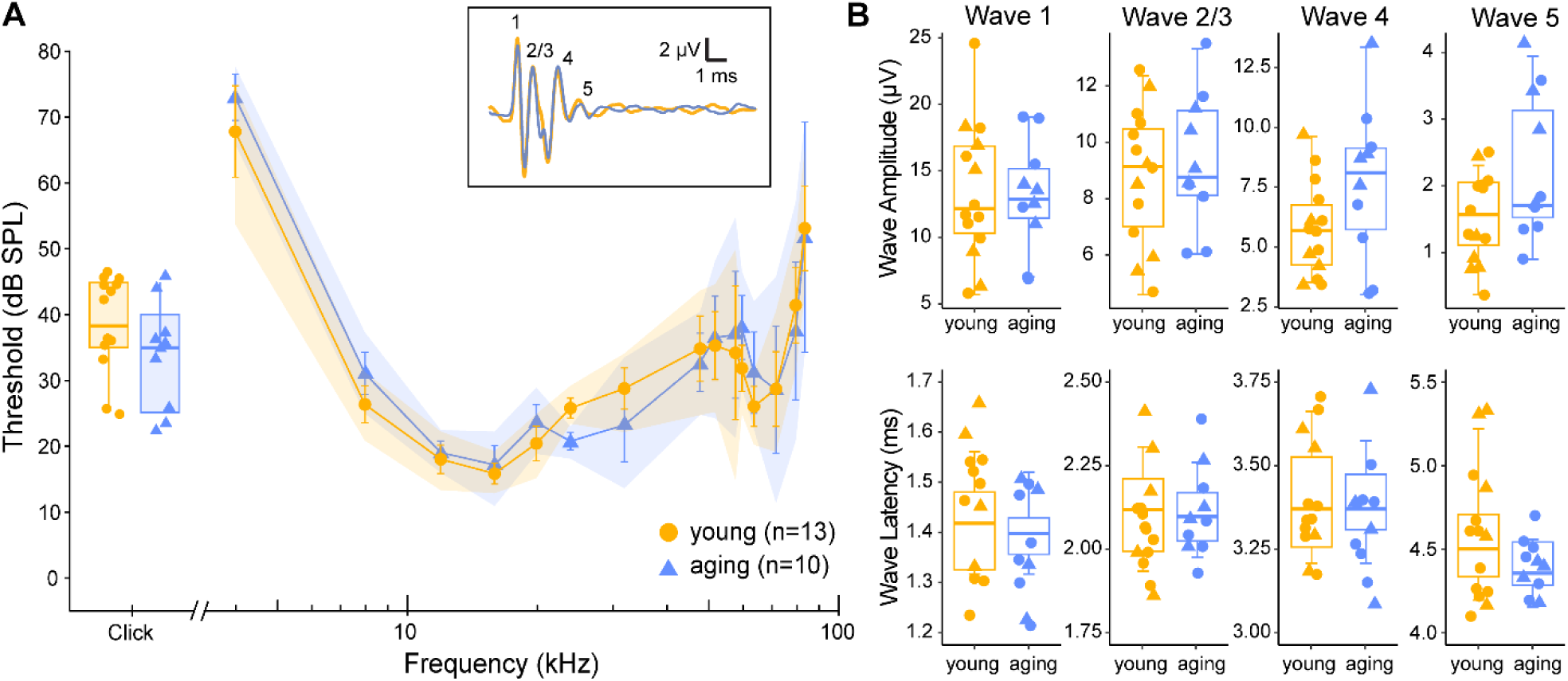
Hearing sensitivity in young and aging *E. fuscus*. **A)** Mean ABR-derived detection thresholds evoked by broadband clicks are shown as boxplots. Curves and error bars represent mean tone-evoked thresholds ± standard error of the mean; 95% confidence intervals are indicated as shaded ribbons for each audiogram. Grand mean ABR waveforms (inset) show similar morphology across age groups. **B)** Peak-to-trough amplitudes (top panels) and latencies (bottom panels) for the individual waves 1-5 of the click-evoked ABR are represented as boxplots with individual data points for young and aging, male (triangles) and female (circles) *E. fuscus*.

The ABR in *E. fuscus* is characterized by 4-5 biphasic waves (Figure 1A, inset panel), following the typical mammalian pattern. In young *E. fuscus*, ABR wave 1, representing activity in the auditory nerve, was typically evoked within approximately 1.4 ms of stimulation (Supplemental Table S1). Waves 2 and 3, representing early brainstem auditory processing centers, often merged to form a single wave approximately 2.1 ms following stimulus onset. Waves 4 and 5, representing higher-order brainstem and midbrain nuclei, occurred within 3.4 and 4.6 ms of stimulus onset, respectively.

The first wave of the ABR is commonly used as an indicator of cochlear integrity, as reduced amplitude and increased latency of ABR wave 1 can reflect early neural contributors to ARHL (e.g., cochlear synaptopathy) and may precede threshold elevation [12,15,16,47]. Wave 1 amplitudes of the click-evoked ABR showed no significant effects of age group (F_1,20.22_=0.18, p=0.68), sex (F_1,20.22_=0.07, p=0.79), or their interaction (F_1,20.22_=0.67, p=0.42) – Supplemental Table S2. There was a significant interaction of age group across stimulus levels on wave 1 amplitudes (F_3,63.25_=4.93, p=3.9×10^-3^); however, post-hoc analyses showed no significant pairwise differences (Supplemental Table S3). Waves 2/3 through 5 of the click-evoked ABR tended to be of greater amplitude in aging bats (Figure 1B) with group differences ranging from 0.47 µV for wave 2/3 to 1.76 µV for wave 4. The amplitudes of ABR waves 4 and 5 showed a significant interaction effect of age and sex (wave 4: F_1,20.38_=5.26, p=0.03; wave 5: F_1,20.31_=4.82, p=0.04) in which aging male bats had significantly larger wave amplitudes relative to young males (wave 4: p=7.0×10^-3^, wave 5: p=0.02).

Click-evoked ABRs had slightly faster wave latencies on average in aging bats relative to young bats (Figure 1B) but these group differences were small, averaging approximately 0.02 for waves 1-4 and 0.20 ms for wave 5. Despite this, wave 4 showed a significant interaction effect of age group and sex on latency (F_1,20.35_=8.61, p=8.1×10^-3^; see Supplemental Table S2 for statistical results for other waves) in which aging male bats showed faster wave latencies relative to younger males (p=0.02). Wave 5 onset occurred earlier in aging males compared to young (Figure 1B), but this trend was not significant, likely owing to variation in wave 5 latency among young males.

We additionally explored age-related changes to tone evoked ABRs for behaviorally relevant frequencies contained within the *E. fuscus* echolocation call: 20-48 kHz corresponding to the spectral content of the fundamental sweep. Although aging bats had slightly larger wave 1 amplitudes evoked by 20-48 kHz, tone-evoked wave 1 amplitudes were similar across age groups (Supplemental Table S4). We observed no significant effects of age group or the interactions of age group with level and age group with sex on the amplitudes or latencies of ABR waves 1-5 for these frequencies (see Supplemental Tables S4-5).

Because within-group variation of ABRs could potentially overshadow more subtle age-related trends, we assessed the relationship of ABR wave 1 amplitude and latency with chronological age using linear regression models fitted to the data. Wave 1 amplitudes showed a slight negative correlation with advancing age for clicks (Figure 2A, 90 dB click: r = -0.05), but not for tones (Figure 2B-C, 24 kHz at 90 dB: r = 0.01, 32 kHz at 90 dB: r = 0.08). Wave 1 latencies were positively correlated with age for 32 kHz tones presented at 90 dB (r = 0.15) indicating slightly delayed responses among aging bats relative to young bats. This trend reversed for lower-level 32 kHz stimuli (from 60-70 dB), in which latencies reduced slightly with age (Figure 2F, Supplemental Table S6). However, there was no significant correlation between age and wave 1 amplitudes or latencies evoked by clicks or tones from 20-48 kHz (Supplemental Table S6), indicating that age-related changes to the ABR were negligible in *E. fuscus*.

**Figure 2.**
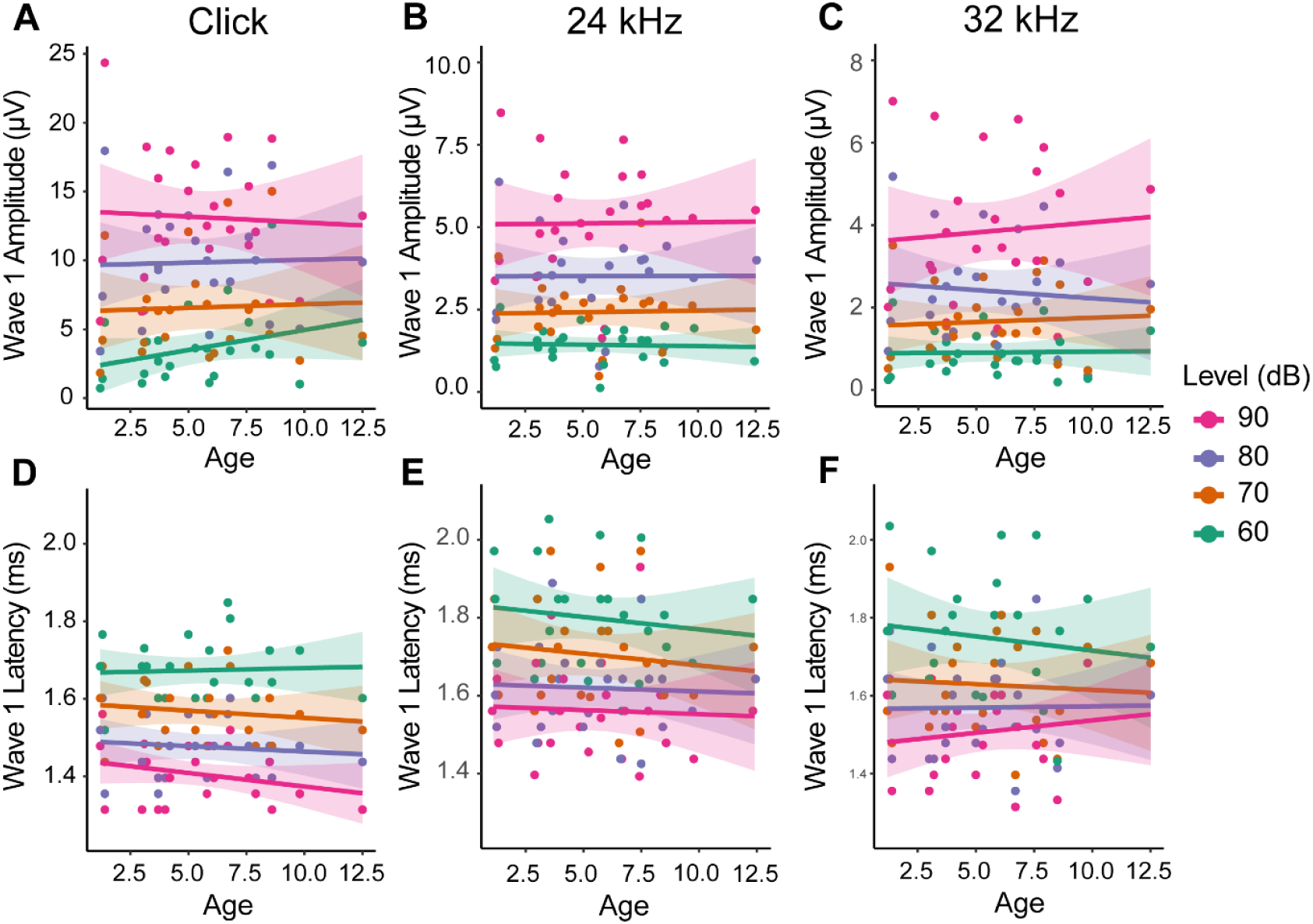
Age effects on *E. fuscus* ABR wave 1 amplitudes (top) and latencies (bottom) evoked by clicks (**A, D**), 24-kHz (**B, E**), and 32-kHz (**C, F**) tones presented from 60-90 dB SPL. Lines represent the linear regression of ABR wave amplitudes across the range of tested ages for each stimulus level; shaded ribbons represent 95% confidence intervals for each linear model.

### Outer hair cell functionality is similar among young and aging bats

To evaluate the functional integrity of the outer hair cells in young and aging bats, we assessed level-dependent changes to the amplitude of the distortion product otoacoustic emission (DPOAE) evoked by *f_2_* frequencies from 8-32 kHz. In young *E. fuscus*, the DPOAE input-output (I/O) function showed a monotonic increase in amplitude per 10 dB increase in stimulus level with a slope approximating 1 (Figure 3). Aging bats showed comparable DPOAE I/O functions to young bats across the frequencies tested, with no significant effects of age group (F_1,180.12_=1.30, p=0.26) or the interactions of age group with frequency (F_1,177.01_=0.46, p=0.50) or age group with level (F_1,177.00_=0.62, p=0.43) on DPOAE amplitudes. DPOAE I/O functions were shifted upwards in aging bats for *f_2_*= 16 and 24 kHz (Figure 3), indicating level-independent increases to amplitudes of otoacoustic emissions relative to young bats, although these differences were not significant.

**Figure 3.**
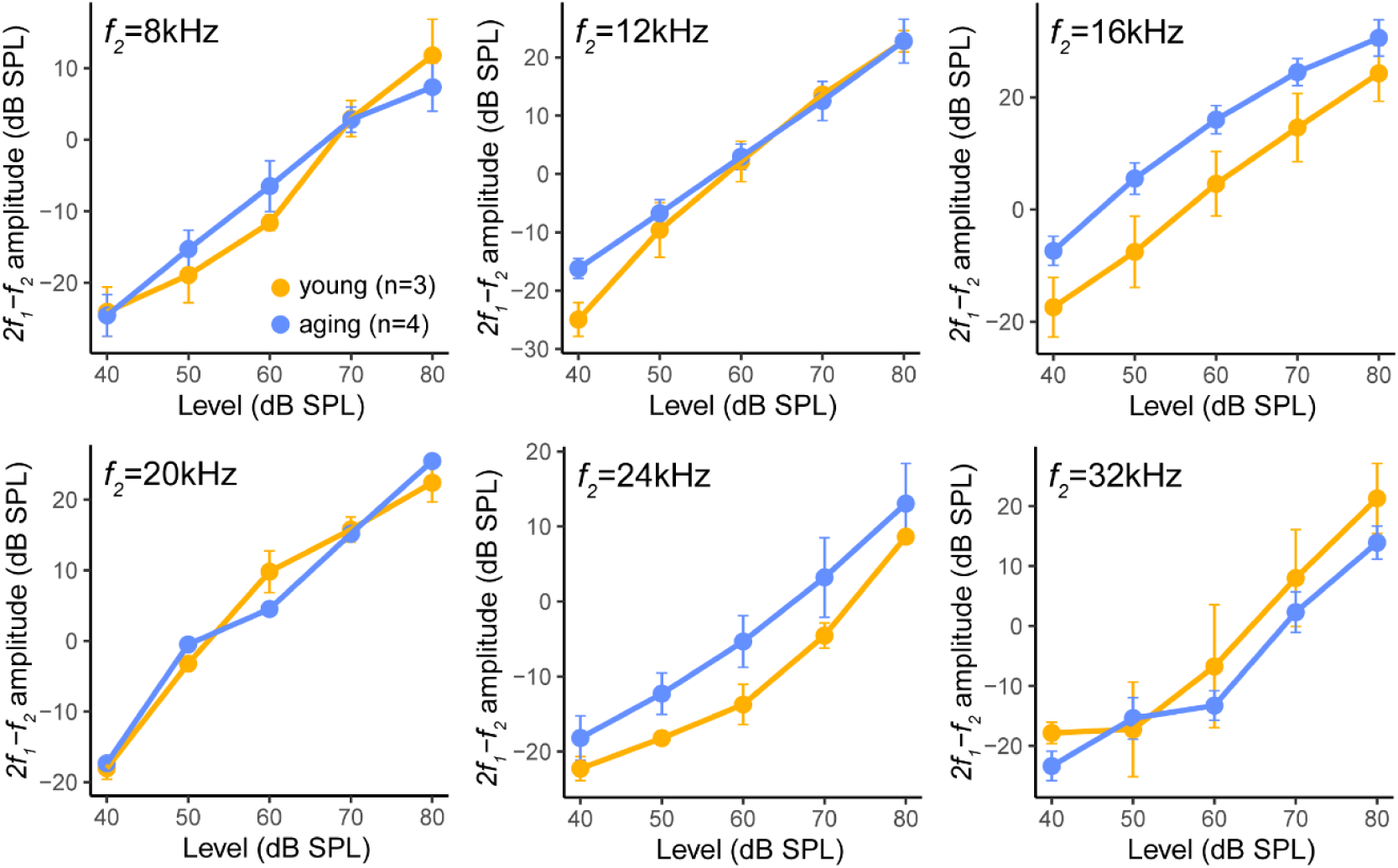
DPOAE amplitudes as a function of stimulus level in young and aging *E. fuscus.* DPOAE input-output curves represent mean amplitudes of emissions evoked by stimulus frequencies *f_2_*: 8-32 kHz presented from 20-80 dB SPL; error bars show standard error of the mean.

### Bat cochlear morphology shows minimal effects of aging

Age-related changes to structural morphology of the cochlea were assessed using whole-mount immunofluorescent preparations from young and aging *E. fuscus* (Figure 4A-B). The aging bat cochlea showed evidence for a slight, non-significant reduction in the number of inner hair cells along the length of the cochlea compared to young bats (Figure 4C, F_1,127.48_=0.61, p=0.44). We observed no age-related variance in the number of presynaptic ribbons in *E. fuscus* (Figure 4D, F_1,53.14_=0.05, p=0.83), with young and aging bats averaging 27-29 ribbons per inner hair cell. However, the overall size of the ribbons, quantified as the total area of CtBP2-positive immunopuncta per inner hair cell, was generally smaller in the aging bat cochlea (8.6 µm^2^ per cell) compared to young (12.1 µm^2^ per cell). Presynaptic ribbons were smallest at locations closer to the apex and base of the aging bat cochlea (Figure 4E), but ribbon size was not significantly affected by age group (F_1,15.13_=3.99, p=0.06) or the interaction of age group and location along the cochlea (F_1,53.08_=0.57, p=0.45).

**Figure 4.**
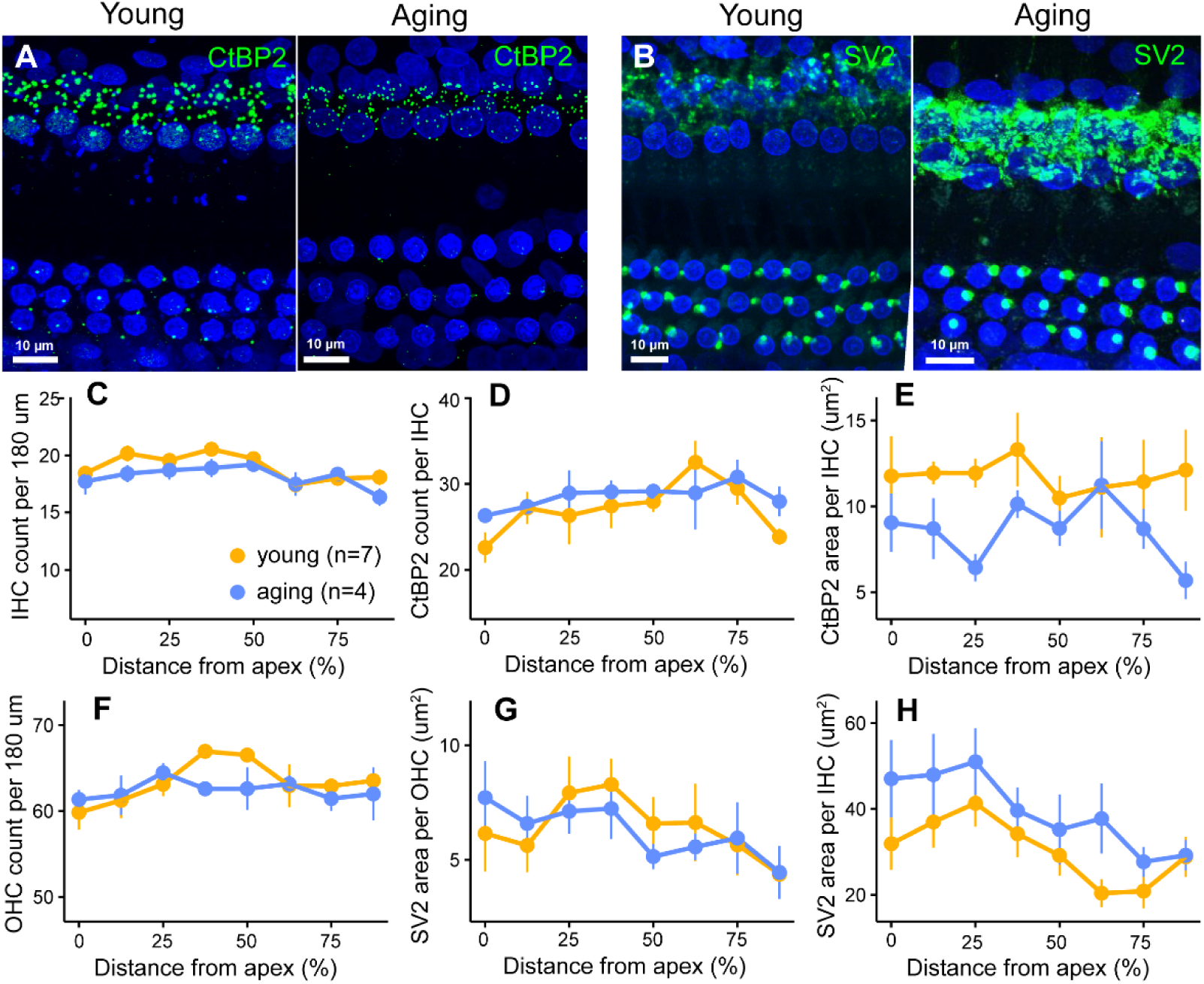
Cochlear immunolabeling in young and aging *E. fuscus*. Maximum projections of the middle turn of the cochlea from a young (1.4-year-old) and aging (12.5-year-old) *E. fuscus.* Cochlear tissue is labeled with cellular marker DAPI (blue), (**A)** presynaptic ribbon marker CtBP2 (green), and **(B)** efferent terminal marker SV2 (green). **C)** Inner hair cell counts in young and aging bats, along with **D, E)** quantification of presynaptic CtBP2-positive immunopuncta per inner hair cell. **F)** Outer hair cell counts in young and aging bats, with quantification of SV2-positive efferent terminals per **G)** outer and **H)** inner hair cell. **C-H:** Data represent mean structural measurements (± standard error) quantified from 8 equidistant regions measured as percent total distance from the apex.

Additionally, aging bats showed a slight reduction in number of outer hair cells at the middle turn of the cochlea approximately 50-62.5% distance from the apex compared to young bats (Figure 4F) but, as with the inner hair cells, this age difference was not significant (F_1,126.64_=2.23, p=0.14). Efferent innervation of the outer hair cells, quantified as total area of SV-2 positive immunopuncta per cell, were comparable across age groups along the length of the cochlea (Figure 4G) indicating no significant age-related changes to the medial olivocochlear system in big brown bats (F_1,65.10_=0.11, p=0.74). In contrast, the density of efferent innervation of the inner hair cells, representing the lateral olivocochlear terminals, appeared to be enhanced among aging bats, particularly at the apical, low-frequency regions of the cochlea (Figure 4H). Despite this, the area of SV2-positive efferent terminals per inner hair cell was not significantly different across age groups (F_1,17.29_=0.32, p=0.58), nor was there an effect of the interaction of age group and location along the cochlea (F_1,64.16_=0.24, p=0.62).

## Discussion

### Aging big brown bats retain ‘youthful’ auditory sensitivity

Hearing is one of the most important sensory modalities for echolocating bats, as the ability to detect low intensity returning echoes is essential for biosonar. Echolocating bats are auditory specialists that are exceptionally long-lived for their size, therefore the selective pressures that shape longevity and echolocation may also drive the retention of sufficient hearing capabilities into old age. The typical trajectory of ARHL in most mammals, including humans, is a progressive loss of sensitivity to sound beginning with high frequency deficits that extend to low frequencies over time [9–11,48]. In the present study we investigated the effects of aging on peripheral auditory sensitivity and cochlear structural integrity in an auditory specialist species, the echolocating big brown bat (*Eptesicus fuscus*). We showed that aging big brown bats retain comparable peripheral hearing sensitivity to young, demonstrated by similar ABR-derived hearing thresholds in animals ranging from 1.2-12.5 years old. The ABR-derived audiogram measured in aging bats overlapped with the young bat audiogram, extending across the same range of frequencies, with peak sensitivity from 12-24 kHz and 60-72 kHz. In contrast, mouse models that show ‘normal’ (i.e., not accelerated) ARHL typically exhibit the onset of behavioral and physiological sensitivity deficits at 15 months that becomes severe (threshold elevation of >60 dB) around 18 months of age (e.g., upon reaching 50% of their maximum 36 month lifespan) [49,50]. We did not observe similar levels of hearing loss in aging big brown bats, indicating that this long-lived species is resistant to, or exhibits considerably delayed, age-related hearing loss.

Additionally, we found that the first wave of the ABR, representing the auditory nerve response and typically considered a metric for cochlear functional integrity, showed minimal non-significant amplitude reductions across age groups. Cochlear aging in humans is correlated with a progressive decline in hair cells, spiral ganglion cells, and auditory nerve fibers [9,15,51], a phenomenon that is mirrored in animal models for ARHL that show degeneration of hair cells, ribbon synapses, and spiral ganglion cells with advancing age [16,47,49,52]. These structural changes are reflected physiologically as reductions to suprathreshold amplitudes of ABR wave 1, which may precede evidence for threshold elevation [16]. In rodents, age-related reduction to ABR wave amplitudes can be profound, with aged animals commonly showing more than 65% reduction of wave 1 of the click-evoked response by 18 months of age [16,49,53,54]. In contrast, suprathreshold wave 1 amplitudes appeared robust to aging in big brown bats, with comparatively small (approximately 11%) decrements to click-evoked wave 1 amplitudes observed in the oldest bats. Further, there were no significant age-related differences in wave 1 amplitudes evoked by frequencies characteristic of the species-specific echolocation call (20-48 kHz, matching spectral content of the fundamental sweep [55]), indicating that aging big brown bats may be resistant to auditory functional declines typically associated with natural senescence.

The later waves of the ABR, waves 4 and 5, showed evidence for sex-biased amplitude enhancements with age in which wave 4 increased in amplitude by approximately 160% and wave 5 more than doubled in aging male bats compared to young males. This change in waveform morphology was accompanied by a non-significant decrease in ABR wave 1 of approximately 0.5 µV (compared to the 0.16 µV decrement in aging female bats that did not show late wave enhancements of such high magnitude). In aging mice and humans, reductions to ABR wave 1 are correlated with stable or increased amplitudes of the later waves that suggest central gain as a compensatory response to age-related deterioration of cochlear input [13,16,56–59]. Although the reduction to wave 1 in *E. fuscus* was not statistically significant and minimal relative to wave 1 reductions observed in rodent models, even a small modification to peripheral hearing capability may be biologically significant to echolocating species that require acute sensitivity to detect low-intensity echoes. The observed changes to ABR wave morphology in aging male bats could represent similar compensatory activity in higher-order brainstem and midbrain auditory processing centers to maintain sensitivity to acoustic signals despite age-related reductions to the cochlear response.

Healthy cochlear function is characterized by compressive, nonlinear amplification driven by the active electro-motile properties of the outer hair cells, which confers greater sensitivity and sharper frequency tuning [60,61]. Outer hair cell loss often precedes inner hair cell loss in the aging mammalian cochlea, which can reduce auditory sensitivity independent of inner hair cell loss [9,49,62]. Age-related changes to outer hair cell function in humans and rodent models for hearing loss is correlated with reduced DPOAEs and shallow, right-shifted growth functions indicating reduced efficacy of the cochlear amplifier [63–65]. In contrast, outer hair cell function in *E. fuscus* was unaffected by age, with no evidence for significant changes to DPOAE amplitudes across stimulus frequency or level. Comparable DPOAE growth functions for frequencies from 8-32 kHz in young and aging bats further indicate that *E. fuscus* maintains robust cochlear response properties into old age which may support continued sensitivity to echolocation signals.

### Bat cochlear structures show signs of noise exposure, but not aging

The peripheral auditory system in mammals is especially susceptible to the cumulative effects of natural senescence and noise damage over a lifetime of exposure to sound. ARHL is commonly associated with the loss of cochlear hair cells, with outer hair cells being particularly susceptible [9,49,51,62]; however, mounting evidence suggests that the perceptual deficits associated with ARHL (e.g., threshold elevation, difficulty listening in noise) are associated with degeneration of the synaptic interface between inner hair cells and auditory afferent fibers [12,13,16,47]. Aging CBA/CaJ mice, a strain used to model a normal, human-like trajectory of ARHL, show up to 50% loss of inner hair cells from the high-frequency basal region of the cochlea and up to 70% loss of outer hair cells towards the low-frequency apical region [49]. Additionally, aging mice show up to 50% loss of ribbon synapses prior to showing evidence for hair cell loss and deafferentation [12,16,66,67].

In the present study, we found no significant evidence for age-related loss of inner or outer hair cells in big brown bats. Further, we observed no significant changes to the number of presynaptic ribbons mediating the afferent auditory pathway in aging bats. However, ribbons were slightly reduced in size in aging bats, providing some potential evidence of senescent changes to cochlear structures. In contrast, synaptopathy in the aging mouse cochlea is associated with enlargement of the remaining inner hair cell ribbons, perhaps as a compensatory mechanism for reduced sensory inputs but also potentially contributing to maladaptive hyperacusis-like responses [67–69]. In aging big brown bats, small ribbon sizes may reflect the consequences of a history of exposure to loud sounds, which has been correlated with reductions to ribbon size in guinea pigs [70].

Although there were no significant age-related differences in efferent innervation of the outer hair cells by medial olivocochlear (OC) neurons, we observed a trend for slightly increased efferent innervation of inner hair cells by lateral OC neurons in aging bats. The OC system enhances auditory detection of signals in noise and may confer protection against acoustic injury via top-down control of the cochlear response (reviewed in [71,72]). The lateral OC system is composed of unmyelinated neurons that originate in and around the lateral superior olive and terminate on the type I afferent fibers that synapse with inner hair cells [71]. Compared to the medial OC, little is known about the mechanism of action of the lateral OC system [73]; however, the lateral OC is hypothesized to prevent acoustic overexposure by protecting against excitotoxicity at the IHC-afferent synapse [74–77]. A greater density of lateral OC efferent terminals per inner hair cell in aging bats could be another structural indicator of a history of noise exposure suggestive of enhanced efferent modulation to prevent noise-induced cochlear damage.

### Echolocating bats as an informative model for resistance to aging and noise

Echolocating bats represent a powerful research model for investigating mechanisms that may protect against noise- and age-related auditory damage: naturally behaving bats regularly emit intense biosonar sounds that subject the auditory system to potentially damaging self-generated sound while simultaneously requiring acute sensitivity to detect low-intensity returning echoes. Hearing loss could prove fatal to an echolocating bat in the wild, as the inability to detect quiet echo returns would prevent effective foraging, navigation, and obstacle avoidance. Although echolocation and intense sound exposure are inextricably linked in bats, the combined physiological and histological results of this study suggest that there is no functional loss of hearing in aging big brown bats.

A limitation of this study is that we were unable to sample bats at the higher end of the maximum longevity for this species (e.g., bats nearing 19 years of age). Therefore, it is possible that we were not able to fully characterize auditory changes that may occur in this species at very old ages. Despite this, we demonstrate that aging big brown bats retain ‘youthful’ physiological hearing thresholds and frequency ranges of sensitivity for up to 12.5 years of life. In contrast, a recent study using Egyptian fruit bats (*Rousettus aegyptiacus*) revealed that not all bats are resistant to ARHL, with individuals showing evidence for high-to-low frequency hearing deficits, comparable to those observed in aging humans [37]. Cross-species variability in susceptibility to ARHL may reflect the different selective pressures experienced by bats, depending on what sensory channels are available to guide behaviors critical for survival. For example, Egyptian fruit bats are able to integrate lingual echolocation with vision during foraging and navigation, with recent evidence suggesting that this species preferentially uses visual over acoustic cues [39]. In contrast, frequency-modulated echoes from self-generated biosonar signals provide the primary sensory cue that enables big brown bats to pursue prey while in flight [78,79]. As such, it is possible that aerial-hawking species like *E. fuscus* possess auditory specializations to protect against hearing loss (reviewed in [80]). Variable susceptibility to noise exposure in echolocating vs. non-echolocating (i.e., visually dominant) bat species presents a complementary view [38], and highlights the value of comparing diverse species to gain a greater understanding of what mechanisms may support cochlear structural and functional integrity in some, but not all, bat species.

In bats, the mechanisms that support exceptional longevity may also play a role in protecting against cochlear damage throughout a lifetime of exposure to sound. ARHL is linked to peripheral sensorineural damage, including loss of cochlear hair cells, loss of afferent and efferent neurons, as well as degeneration of the stria vascularis, an energetically active structure that maintains the endocochlear potential. In particular, the accumulation of reactive oxygen species (ROS) and subsequent ROS-driven mitochondrial dysfunction has been implicated in cochlear aging, including in the deterioration and loss of hair cells and atrophy of the stria [22,81,82]. Many bat species appear to be resistant to oxidative stress [28,33,83,84], an attribute that supports homeostatic processes over a long lifespan and may similarly protect against age-related accumulation of oxidative damage in the cochlea. Despite this, ARHL in the Egyptian fruit bat is correlated with clear structural markers of strial atrophy [37]. Careful investigation of the stria vascularis in young and aging big brown bats is a key next step to evaluate whether this species shows resistance to age-related degeneration of this structure.

Additionally, auditory sensitivity appears robust to acute noise exposures in bats [44–46,80]. The protective factors that may underlie resistance to noise in bats remain unknown; however, the auditory efferent system presents an intriguing avenue for future research. Previous morphological studies have revealed that the efferent OC system is hypertrophic in bats, with some species showing nearly three times the number of efferent OC neurons observed in mice [85–87]. Here we observed similar OC enhancement in *E. fuscus,* with young bats showing twice the area of efferent synapses per inner and outer hair cell compared to young mice [49]. Strong top-down efferent modulation of the cochlear response to sound has been observed in bats living in noisy roosts [88] and during call production [89] as a protective mechanism against self-generated and background sounds. Additionally, the middle ear muscle reflex, which is highly developed in bats and contracts in coordination with the laryngeal muscles during echolocation [90–93], may further protect against the accumulation of cochlear damage from intense self-generated sounds. Further physiological and behavioral investigation of efferent function in young and aging bats, particularly those that utilize active and passive listening in noisy environments, will extend our understanding of the mechanisms that allow these long-lived auditory specialists to resist the damaging effects of intense sounds.

## Methods

### Animals

We assessed auditory sensitivity in 23 wild-caught *E. fuscus* collected in the state of Maryland (MD Dept. of Natural Resources, permit #55440). Bats were housed with conspecifics in a colony with sufficient roosting locations, room to fly, and access to food and water *ad libitum*. Colony housing was maintained at 21-27°C with a relative humidity of 30-70%. All procedures were performed with the approval of the Johns Hopkins Animal Care and Use Committee and complied with the National Institutes of Health guide for the care and use of laboratory animals.

Bats were grouped by age into young (n=13, 8 female, 5 male) and aging (n=10, 6 female, 4 male) categories for analyses. Aging phenotypes, including life stage definitions, have not been established in bats; therefore, we categorized bat age groups based on percent maximum longevity. Maximum longevity for *E. fuscus* is reportedly 19 years [42]; however, adult survivorship is low among wild populations that show a 50% survivorship of 2-3 years [94–96] and mean longevity of approximately 6 years [43]. In the present study, bats younger than 6 years of age (<40% maximum species longevity) were considered young and bats older than 6 years (>40% maximum species longevity) were considered to be aging. We selected age groupings similar to those used in other mammalian aging studies, where ‘old’ phenotypes are typically characterized by the onset of senescent physiological changes in some but not all biomarkers, and ‘very old’ phenotypes display more severe deficits in all aging biomarkers [97]. For example, previous aging studies using short-lived laboratory mice and comparably long-lived deer mice have categorized animals as ‘old’ when age exceeds 40-50% maximum longevity [98–105].

### Age estimation

The laboratory colony is comprised of wild-caught *E. fuscus* that are of known minimum ages based on collection dates. To allow for more precise age estimation of the individuals used in the present study, we used DNA methylation (DNAm) profiling following established procedures described in [106]. DNA was extracted from wing tissue biopsies (3-4mm diameter) and processed using the Zymo Quick-DNA Miniprep Plus Kit (ZymoResearch, Orange, CA). Samples were submitted to the Clock Foundation (Torrance, California), to measure the proportion of methylated and unmethylated sites using a custom Illumina microarray [107]. We estimated ages using a species-specific epigenetic clock generated using 60 *E. fuscus* bats of known age [106]. This DNA methylation procedure has been previously validated to provide estimates of chronological age with high accuracy [106].

### Auditory brainstem response recording

We recorded auditory brainstem responses (ABRs) following previously published procedures [36]. Briefly, bats were anesthetized via an intraperitoneal injection of 50 mg/kg ketamine and 30 mg/kg xylazine and placed on a 37°C warming pad in a sound attenuating chamber lined with acoustic foam (IAC Acoustics). Auditory evoked potentials were recorded using three subdermal needle electrodes placed at the vertex of the skull (recording), along the mastoid (inverting) and in the shoulder (ground). ABRs were acquired using BioSigRZ software (Tucker Davis Technologies (TDT), Alachua, FL) at a 12 kHz sampling rate with a TDT Medusa 4Z preamplifier connected to a TDT RZ6 I/O processor.

Sound-evoked potentials were recorded in response to broadband clicks and tones from 4-84 kHz (5 ms tone pips with 0.5 ms cos^2^ on/off ramp) that were generated using TDT SigGenRZ software. Acoustic stimuli were broadcast using free-field speakers (TDT MF1 magnetic speaker for clicks and frequencies below 60 kHz, TDT ES1 electrostatic speaker for frequencies 60 kHz and greater) placed 10 cm from the bat’s head and oriented to present sound directly to the ear along the longitudinal axis of the pinna. ABR stimuli were calibrated to 90 dB peak equivalent SPL using a calibrated ¼ inch microphone (PCB Piezotronics, model 377C01) and were attenuated in 10 dB steps to present a level series from 90-10 dB SPL. Stimuli were presented at a rate of 21/second. ABR waveforms were averaged across 512 stimulus presentations and filtered from 0.3-3 kHz for subsequent analyses.

Detection thresholds were defined as the intermediate stimulus presentation level above which an evoked response was discriminable from the noise floor of the recording and below which no response was observed, following published methods (e.g., [102,108–111]). Physiological recordings were collected prior to completion of age estimation procedures and detection thresholds were determined using visual inspection by two independent, experienced observers blinded to the age group of each subject. ABR wave peak-to-trough amplitudes and latencies were extracted using an automated, user-supervised software [112].

We statistically assessed age effects on ABR detection thresholds and the amplitudes and latencies of individual ABR waves using linear mixed-effects models (LMM) fit with restricted maximum likelihood estimation. We compared detection thresholds across the fixed effects of stimulus frequency, age group, sex, and the interaction of frequency, age group, and sex. We assessed age group differences in ABR wave amplitudes and latencies for click and tone evoked responses across the fixed effects of stimulus level, age group, sex, and the interactions of level, age group, and sex. Wave amplitude data were log-transformed to meet assumptions of the LMMs. We removed nonsignificant interaction terms to reduce model complexity. For all linear mixed-effects models, we included subject ID as a random effect to account for individual variation. We assessed the significance of fixed effects using a conditional F-test with Kenward-Rogers correction for degrees of freedom. Post hoc analyses of significant effects were performed using a Tukey HSD correction for multiple comparisons. We additionally evaluated the relationship between ABR wave amplitudes and latencies and chronological age as a continuous predictor using linear regression followed by hypothesis testing of the Pearson correlation coefficient (r) against the null hypothesis predicting no correlation (r=0) of age and wave amplitude. All statistical testing was performed in R statistical software v.4.3.2 [113].

### Distortion product otoacoustic emission recording

We assessed outer hair cell function in a subset of bats (young n=3, aging n=4) using distortion product otoacoustic emissions (DPOAEs). DPOAEs were recorded using BioSigRZ (TDT) in anesthetized animals within a sound-attenuating chamber, as described in the ABR recording procedures above. Closed-field acoustic stimuli were two simultaneous, iso-intensity tones (*f*_1_ and *f*_2_, *f*_2_ frequency range: 8-32 kHz and *f*_1_ presented at an *f_2_/f_1_* frequency ratio of 1.2) presented by two speakers (TDT MF1) coupled to an ear insert that was tightly inserted into the ear canal. Both speakers were calibrated in-ear for each subject to account for potential variability in ear canal characteristics. Stimuli were presented from 80-20 dB SPL in 10 dB decrements. DPOAEs (*2f_1_-f_2_*) were recorded using a low-noise probe microphone (TDT DPM1, flat frequency response 3-40 kHz) coupled to the ear insert and were averaged over 512 stimulus presentations. We additionally recorded control DPOAEs in a dead bat to confirm that the data collected in living bats reflected active physiological processes rather than distortions generated as stimulus artifacts. We extracted DPOAE amplitudes per stimulus level to generate input-output functions and compared these across age groups using linear mixed-effects modeling in R, incorporating subject ID as a random effect in the model.

### Cochlear immunolabeling

We performed immunohistochemistry to label whole mount cochlear dissections in a subset of bats (young n=7, aging n=4) for a qualitative assessment of cochlear structures across age groups. When possible, bats were transcardially perfused with 4% paraformaldehyde prior to harvesting left and right cochleas; however, in some cases cochleas were obtained post-mortem. Cochleas were post-fixed in 4% paraformaldehyde at 4°C for at least 24 hours prior to decalcification using 0.5M EDTA. Cochleas were then placed into blocking buffer solution (5% normal goat serum, 10% bovine serum albumen, and 0.5% Triton X-100 (Electron Microscopy Services)) for one hour, followed by 24-hour incubation in primary antibodies diluted in blocking buffer. Primary antibodies included anti-myosin 6 (Myo-6, 1:500, Bioss cat# bs-11264R) to label cochlear hair cells, anti-synaptic vesicle protein (SV2, 1:500, DSHB cat# SV2, RRID: AB_2315387) to label efferent olivocochlear synapses, and anti-C-terminal binding protein (CtBP2, 1:200, BD Transduction Laboratories, cat# 612044, RRID: AB_399431) to label presynaptic ribbons. Cochleas were washed three times in phosphate buffered saline (PBS) and transferred to secondary antibodies in blocking buffer for two hours at room temperature. Secondary antibodies used were Alexa Fluor 488 goat anti-mouse SFX (Molecular Probes cat # A31619) and Alexa Fluor 568 goat anti-rabbit IgG (Invitrogen cat# A11036).

Cochleas were dissected into 4-5 flat turns and mounted on slides using DAPI-Fluoromount-G mounting medium (Southern Biotech) for imaging. Whole-mount cochlea preparations were imaged at 10X using a Nikon Eclipse Ti2 confocal microscope. Images were exported and the Measure_Line plugin for ImageJ (Eaton-Peabody Laboratories, Massachusetts Eye and Ear) was used to identify 8 equidistant locations along the cochlea based on percent total length. Confocal z-stacks (z=0.15 µm) of these locations along the cochlea were generated using a 60X/1.4 NA oil-immersion lens. Cochlear structures were analyzed using FIJI [114]. Inner and outer hair cells were quantified per 180 µm continuous segment of each region. Counts and surface area measurements of CtBP2-positive synaptic ribbon puncta and SV2-positive efferent terminals were automatically extracted using the Analyze Particles function in FIJI. We analyzed age-related differences in cochlear structures using linear mixed-effects modeling in R, including age group and cochlear location as fixed effects and subject ID as a random effect.

## Acknowledgements

This work was supported by the following grants: NIH NIDCD T32 DC000023 (G.C., C.A.D.), the David M. Rubenstein Fund for Hearing Research (A.M.L., C.F.M.), NIH NINDS R01 NS121413, NIH NINDS R34 NS118462-01, Office of Naval Research N00014-17-1-2736, Office of Naval Research N00014-22-1-2793, NSF NCS-FO 1734744 (C.F.M.), and NSF DBI-2213824 (G.S.W).

