## Supplemental Material for "Resistance to age-related hearing loss in the echolocating big brown bat (*Eptesicus fuscus*)"

**Supplemental Figures and Tables**

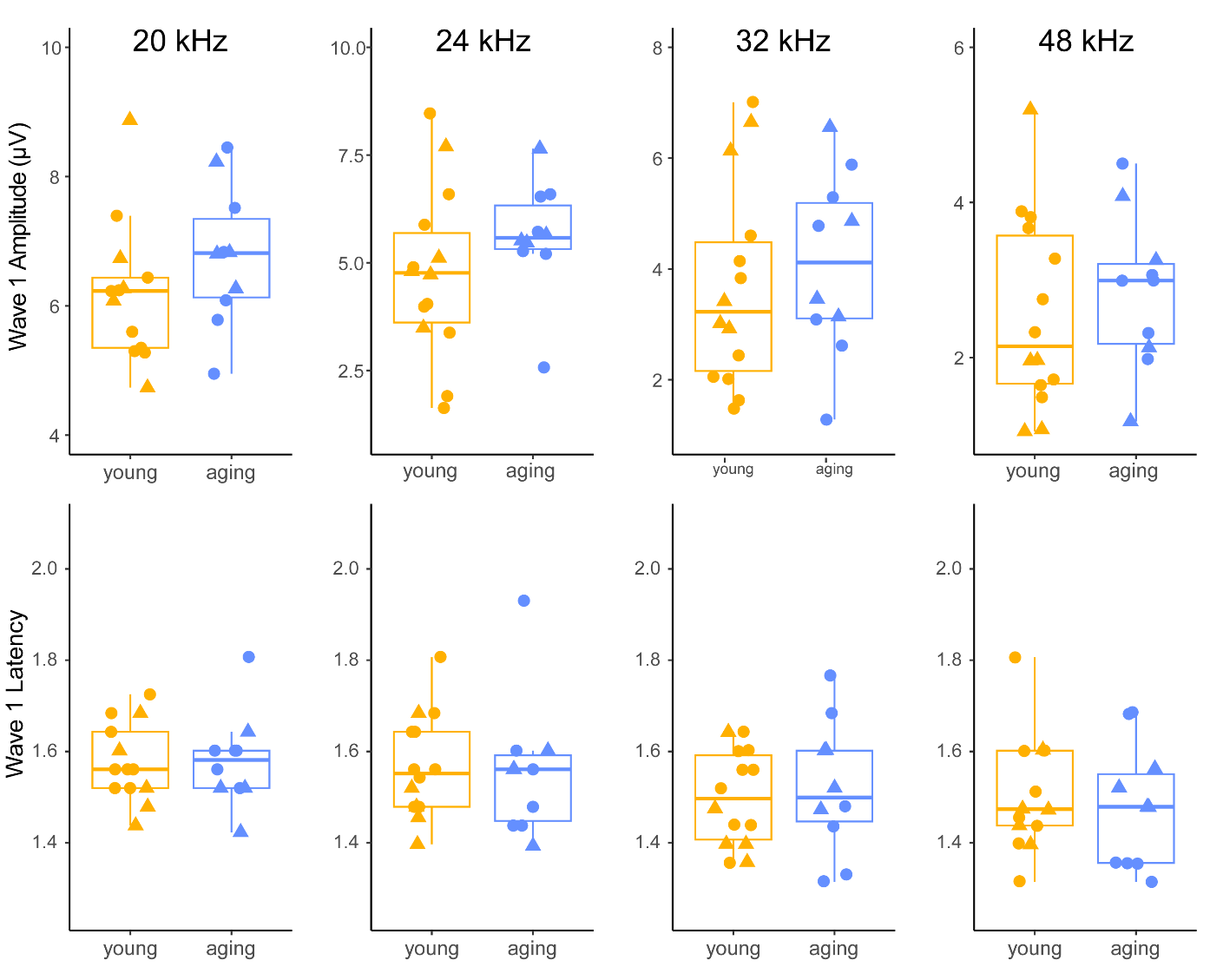
**Figure S1.** Tone-evoked ABR wave 1 metrics in young and aging *E. fuscus*. Peak-to-trough amplitudes (top panels) and latencies (bottom panels) of wave 1 evoked by 20, 24, 32, and 48 kHz tones. Data points indicate male (triangles) and female (circles) *E. fuscus*.

**Table S1.** Average (± SD) ABR wave metrics for responses evoked by 90 dB click stimuli

|  | **Wave** | **Young** | | **Aging** | |
| --- | --- | --- | --- | --- | --- |
|  |  | Female (n=8) | Male (n=5) | Female (n=6) | Male (n=4) |
| **Amplitude**  **(µV)** | 1 | 13.36 (± 5.41) | 13.06 (± 5.25) | 13.19 (± 5.45) | 12.63 (± 1.23) |
|  | 2/3 | 9.18 (± 2.45) | 8.30 (± 2.66) | 9.03 (± 3.03) | 9.82 (± 1.38) |
|  | 4 | 5.98 (± 1.69) | 5.76 (± 2.40) | 6.35 (± 2.95) | 9.62 (± 2.53) |
|  | 5 | 1.81 (± 0.62) | 1.28 (± 0.65) | 1.92 (± 1.02) | 3.01 (± 1.03) |
| **Latency**  **(ms)** | 1 | 1.40 (± 0.09) | 1.43 (± 0.07) | 1.39 (± 0.06) | 1.41 (± 0.08) |
|  | 2/3 | 2.08 (± 0.11) | 2.15 (± 0.13) | 2.11 (± 0.11) | 2.08 (± 0.10) |
|  | 4 | 3.34 (± 0.12) | 3.49 (± 0.18) | 3.42 (± 0.10) | 3.32 (± 0.16) |
|  | 5 | 4.44 (± 0.18) | 4.83 (± 0.45) | 4.42 (± 0.13) | 4.32 (± 0.17) |

**Table S2.** Effects of age and stimulus level on click-evoked ABR wave amplitudes and latencies.

|  | **Factor** | **Sum Sq** | **Mean Sq** | **df** | **F-value** | **P-value** |
| --- | --- | --- | --- | --- | --- | --- |
| *Amplitude* | | | | | | |
| Wave 1 | Age group | 0.009 | 0.009 | 1,20.2 | 0.177 | 0.678 |
|  | Sex | 0.004 | 0.004 | 1,20.2 | 0.072 | 0.791 |
|  | **Level** | **25.31** | **8.438** | **3,63.2** | **174.0** | **<2.2x10^-16^** |
|  | Age group x Sex | 0.033 | 0.033 | 1,20.2 | 0.670 | 0.423 |
|  | **Age group x Level** | **0.717** | **0.239** | **3,63.2** | **4.909** | **0.004** |
| Wave 2/3 | Age group | 0.062 | 0.062 | 1,20.4 | 1.451 | 0.243 |
|  | Sex | 0.026 | 0.026 | 1,20.4 | 0.604 | 0.446 |
|  | **Level** | **27.90** | **9.299** | **3,63.5** | **218.9** | **<2.0x10^-16^** |
|  | Age group x Sex | 0.038 | 0.038 | 1,20.4 | 0.905 | 0.353 |
|  | Age group x Level | 0.106 | 0.035 | 3,63.5 | 0.827 | 0.483 |
| Wave 4 | **Age group** | **0.315** | **0.315** | **1,20.4** | **5.982** | **0.024** |
|  | Sex | 3.0x10^-4^ | 3.0x10^-4^ | 1,20.4 | 0.005 | 0.944 |
|  | **Level** | **20.79** | **6.929** | **3,63.5** | **131.5** | **<2.0x10^-16^** |
|  | **Age group x Sex** | **0.277** | **0.277** | **1,20.4** | **5.258** | **0.033** |
|  | Age group x Level | 0.038 | 0.013 | 3,63.5 | 0.242 | 0.867 |
| Wave 5 | Age group | 0.773 | 0.773 | 1,20.3 | 2.928 | 0.102 |
|  | Sex | 0.051 | 0.051 | 1,20.3 | 0.194 | 0.665 |
|  | **Level** | **15.18** | 5.060 | **3,63.9** | **19.17** | **5.6x10^-9^** |
|  | **Age group x Sex** | **1.272** | **1.272** | **1,20.3** | **4.820** | **0.040** |
|  | Age group x Level | 0.612 | 0.204 | 3,63.9 | 0.773 | 0.513 |
| *Latency* | | | | | | |
| Wave 1 | Age group | 2.2x10^-5^ | 2.2x10^-5^ | 1,20.2 | 2.0x10^-3^ | 0.989 |
|  | Sex | 9.2x10^-4^ | 9.2x10^-4^ | 1,20.2 | 1.138 | 0.299 |
|  | **Level** | **0.907** | **0.302** | **3,63.2** | **374.4** | **<2.0x10^-16^** |
|  | Age group x Sex | 2.9x10^-4^ | 2.9x10^-4^ | **1,20.2** | 0.362 | 0.554 |
|  | *Age group x Level* | *6.3x10^-3^* | *2.1x10^-3^* | 3,63.2 | *2.623* | *0.058* |
| Wave 2/3 | Age group | 1.4x10^-3^ | 1.4x10^-3^ | 1,20.2 | 0.465 | 0.503 |
|  | Sex | 5.5x10^-4^ | 5.5x10^-4^ | 1,20.2 | 0.182 | 0.673 |
|  | **Level** | **1.548** | **0.516** | **3,63.2** | **171.7** | **<2.0x10^-16^** |
|  | Age group x Sex | 3.3x10^-3^ | 3.3x10^-3^ | **1,20.2** | 1.101 | 0.306 |
|  | Age group x Level | 2.5x10^-3^ | 8.2x10^-4^ | 3,63.2 | 0.272 | 0.845 |
| Wave 4 | Age group | 0.009 | 0.009 | 1,20.3 | 1.399 | 0.250 |
|  | Sex | 0.005 | 0.005 | 1,20.3 | 0.793 | 0.384 |
|  | **Level** | **1.967** | **0.655** | **3,63.4** | **101.5** | **<2.2x10^-16^** |
|  | **Age group x Sex** | **0.056** | **0.056** | **1,20.3** | **8.613** | **0.008** |
|  | Age group x Level | 1.0x10^-4^ | 3.4x10^-4^ | 3,63.4 | 0.053 | 0.984 |
| Wave 5 | Age group | 5.1x10^-4^ | 5.1x10^-4^ | 1,20.4 | 0.211 | 0.652 |
|  | Sex | 3.9x10^-4^ | 3.9x10^-4^ | 1,20.4 | 0.171 | 0.692 |
|  | **Level** | **0.333** | **0.111** | **3,63.6** | **45.50** | **7.4x10^-16^** |
|  | Age group x Sex | 0.007 | 0.007 | **1,20.4** | 2.888 | 0.105 |
|  | Age group x Level | 0.014 | 0.005 | 3,63.6 | 1.947 | 0.131 |

Results of linear mixed-effects modeling testing the significance of fixed effects (age group, level, sex, and the interactions of age group and sex and age group and level) using R package lme4. Conditional F-tests (Type III) were performed with Kenward-Roger’s method for degrees of freedom calculation using R package LmerTest.

**Table S3.** Pairwise comparisons of click-evoked ABR wave 1 amplitudes in young and aging bats across stimulus levels.

| **Level** | **Estimate** | **t-ratio** | **P-value** | **Effect size** | **95% CI** |
| --- | --- | --- | --- | --- | --- |
| 60 dB | -0.403 | -1.701 | 0.101 | -1.831 | -4.07, 0.41 |
| 70 dB | 0.021 | 0.086 | 0.932 | 0.093 | -2.12, 2.31 |
| 80 dB | -0.029 | -0.123 | 0.903 | -0.132 | -2.35, 2.08 |
| 90 dB | 0.039 | 0.166 | 0.869 | 0.177 | -2.01, 2.37 |

Results of post-hoc pairwise comparisons for significant fixed effects. P-values calculated using Tukey HSD correction for multiple comparisons using R package emmeans.

**Table S4.** Effects of age and stimulus level on tone-evoked ABR wave amplitudes.

|  |  | **Factor** | **Sum Sq** | **Mean Sq** | **df** | **F-value** | **P-value** |
| --- | --- | --- | --- | --- | --- | --- | --- |
|  | ***20 kHz*** | | | | | | |
|  | Wave 1 | Age group | 0.181 | 0.181 | 1,19 | 3.471 | 0.078 |
|  |  | Sex | 0.095 | 0.095 | 1,19 | 1.813 | 0.194 |
|  |  | **Level** | **38.01** | **12.67** | **3,63** | **243.1** | **<2.0e-16** |
|  |  | Age group x Sex | 0.082 | 0.082 | 1,19 | 1.573 | 0.225 |
|  |  | Age group x Level | 0.313 | 0.104 | 3,63 | 2.002 | 0.123 |
|  | Wave 2/3 | Age group | 0.072 | 0.072 | 1,19 | 0.710 | 0.410 |
|  |  | Sex | 0.022 | 0.022 | 1,19 | 0.220 | 0.644 |
|  |  | **Level** | **47.64** | **15.88** | **3,63** | **157.0** | **<2.0e-16** |
|  |  | Age group x Sex | 0.405 | 0.405 | 1,19 | 4.000 | 0.060 |
|  |  | Age group x Level | 0.729 | 0.243 | 3,63 | 2.403 | 0.076 |
|  | Wave 4 | Age group | 0.003 | 0.003 | 1,19 | 0.050 | 0.825 |
|  |  | Sex | 0.010 | 0.010 | 1,19 | 0.156 | 0.696 |
|  |  | **Level** | **37.67** | **12.56** | **3,63** | **192.6** | **<2.0e-16** |
|  |  | Age group x Sex | 0.044 | 0.044 | 1,19 | 0.668 | 0.421 |
|  |  | Age group x Level | 0.475 | 0.158 | 3,63 | 2.430 | 0.073 |
|  | Wave 5 | Age group | 0.136 | 0.136 | 1,19 | 0.806 | 0.381 |
|  |  | Sex | 0.138 | 0.138 | 1,19 | 0.820 | 0.377 |
|  |  | **Level** | **22.68** | **7.562** | **3,63** | **44.98** | **<1.2e-15** |
|  |  | Age group x Sex | 0.032 | 0.032 | 1,19 | 0.188 | 0.669 |
|  |  | Age group x Level | 0.397 | 0.132 | 3,63 | 0.787 | 0.505 |
| ***24 kHz*** | | | | | | | |
| Wave 1 | | Age group | 0.040 | 0.040 | 1,20 | 1.045 | 0.319 |
|  |  | Sex | 0.046 | 0.046 | 1,20 | 1.183 | 0.290 |
|  |  | **Level** | **21.99** | **7.331** | **3,66** | **189.4** | **<2.0e-16** |
|  |  | Age group x Sex | 0.018 | 0.018 | 1,20 | 0.463 | 0.504 |
|  |  | Age group x Level | 0.007 | 0.002 | 3,66 | 0.063 | 0.979 |
| Wave 2/3 | | Age group | 0.037 | 0.037 | 1,20 | 1.025 | 0.324 |
|  |  | Sex | 0.005 | 0.005 | 1,20 | 0.132 | 0.720 |
|  |  | **Level** | **16.13** | **5.378** | **3,66** | **147.9** | **<2.0e-16** |
|  |  | Age group x Sex | 0.003 | 0.003 | 1,20 | 0.081 | 0.779 |
|  |  | Age group x Level | 0.131 | 0.044 | 3,66 | 1.205 | 0.315 |
| Wave 4 | | Age group | 0.300 | 0.300 | 1,20 | 4.139 | 0.055 |
|  |  | **Sex** | **0.376** | **0.376** | **1,20** | **5.193** | **0.034** |
|  |  | **Level** | 13.70 | 4.565 | **3,66** | 63.03 | **<2.0e-16** |
|  |  | Age group x Sex | 0.008 | 0.008 | 1,20 | 0.110 | 0.744 |
|  |  | Age group x Level | 0.171 | 0.057 | 3,66 | 0.787 | 0.506 |
| Wave 5 | | Age group | 0.330 | 0.330 | 1,20 | 1.623 | 0.217 |
|  |  | Sex | 0.131 | 0.131 | 1,20 | 0.645 | 0.431 |
|  |  | **Level** | **11.68** | **3.894** | **3,66** | **19.16** | **4.8e-09** |
|  |  | Age group x Sex | 0.166 | 0.166 | 1,20 | 0.817 | 0.377 |
|  |  | Age group x Level | 1.025 | 0.342 | 3,66 | 1.680 | 0.180 |
| ***32 kHz*** | | | | | | | |
| Wave 1 | | Age group | 0.000 | 0.000 | 1,20 | 0.001 | 0.983 |
|  |  | Sex | 0.146 | 0.146 | 1,20 | 2.899 | 0.104 |
|  |  | **Level** | **29.00** | **9.662** | **3,66** | **192.0** | **<2.0e-16** |
|  |  | Age group x Sex | 0.008 | 0.008 | 1,20 | 0.162 | 0.692 |
|  |  | Age group x Level | 0.181 | 0.061 | 3,66 | 1.202 | 0.316 |
| Wave 2/3 | | Age group | 0.021 | 0.021 | 1,20 | 0.244 | 0.627 |
|  |  | Sex | 0.051 | 0.051 | 1,20 | 0.609 | 0.444 |
|  |  | **Level** | **21.74** | **7.246** | **3,66** | **86.24** | **<2.0e-16** |
|  |  | Age group x Sex | 0.003 | 0.003 | 1,20 | 0.031 | 0.862 |
|  |  | Age group x Level | 0.198 | 0.066 | 3,66 | 0.786 | 0.506 |
| Wave 4 | | Age group | 0.067 | 0.067 | 1,20 | 0.711 | 0.409 |
|  |  | Sex | 0.159 | 0.159 | 1,20 | 1.691 | 0.208 |
|  |  | **Level** | **14.93** | **4.976** | **3,66** | **53.08** | **<2.0e-16** |
|  |  | Age group x Sex | 0.011 | 0.011 | 1,20 | 0.113 | 0.740 |
|  |  | Age group x Level | 0.277 | 0.092 | 3,66 | 0.986 | 0.406 |
| Wave 5 | | Age group | 0.010 | 0.010 | 1,20 | 0.085 | 0.774 |
|  |  | Sex | 0.025 | 0.025 | 1,20 | 0.206 | 0.655 |
|  |  | **Level** | **7.895** | **2.632** | **3,66** | **21.98** | **5.6e-10** |
|  |  | Age group x Sex | 0.003 | 0.003 | 1,20 | 0.022 | 0.885 |
|  |  | Age group x Level | 0.553 | 0.184 | 3,66 | 1.538 | 0.213 |
|  | ***48 kHz*** | | | | | | |
|  | Wave 1 | Age group | 0.028 | 0.028 | 1,20 | 0.542 | 0.470 |
|  |  | Sex | 0.025 | 0.025 | 1,20 | 0.482 | 0.496 |
|  |  | **Level** | **26.30** | **8.766** | **3,64** | **167.6** | **<2e-16** |
|  |  | Age group x Sex | 0.004 | 0.004 | 1,20 | 0.080 | 0.782 |
|  |  | Age group x Level | 0.073 | 0.024 | 3,64 | 0.464 | 0.707 |
|  | Wave 2/3 | Age group | 0.012 | 0.012 | 1,20 | 0.104 | 0.751 |
|  |  | Sex | 0.262 | 0.262 | 1,20 | 2.233 | 0.151 |
|  |  | **Level** | **20.54** | **6.847** | **3,64** | **58.35** | **<2e-16** |
|  |  | Age group x Sex | 0.043 | 0.042 | 1,20 | 0.362 | 0.554 |
|  |  | Age group x Level | 0.485 | 0.162 | 3,64 | 1.377 | 0.258 |
|  | Wave 4 | Age group | 0.012 | 0.012 | 1,20 | 0.161 | 0.693 |
|  |  | Sex | 0.003 | 0.003 | 1,20 | 0.044 | 0.837 |
|  |  | **Level** | **16.26** | **5.419** | **3,64** | **72.05** | **<2e-16** |
|  |  | Age group x Sex | 0.028 | 0.028 | 1,20 | 0.376 | 0.546 |
|  |  | **Age group x Level** | 0.543 | 0.181 | 3,64 | 2.404 | 0.076 |
|  | Wave 5 | Age group | 0.034 | 0.034 | 1,20 | 0.218 | 0.646 |
|  |  | Sex | 0.387 | 0.387 | 1,20 | 2.455 | 0.133 |
|  |  | **Level** | **9.817** | **3.272** | **3,64** | **20.74** | **1.7e-09** |
|  |  | Age group x Sex | 0.002 | 0.002 | 1,20 | 0.011 | 0.919 |
|  |  | Age group x Level | 0.236 | 0.079 | 3,64 | 0.499 | 0.684 |

Results of linear mixed-effects modeling testing the significance of fixed effects (age group, level, and sex and the interactions of age group and sex and age group and level) on the measured amplitudes of ABRs evoked by 20-48 kHz tones. For all tests, subject ID was included as a random effect to account for individual variation.

**Table S5.** Effects of age and stimulus level on tone-evoked ABR wave latencies.

|  |  | **Factor** | **Sum Sq** | **Mean Sq** | **df** | **F-value** | **P-value** |
| --- | --- | --- | --- | --- | --- | --- | --- |
|  | ***20 kHz*** | | | | | | |
|  | Wave 1 | Age group | 4.5e-4 | 4.5e-4 | 1,19 | 0.358 | 0.557 |
|  |  | Sex | 0.004 | 0.004 | 1,19 | 2.834 | 0.109 |
|  |  | **Level** | **0.497** | **0.166** | **3,63** | **132.4** | **<2.0e-16** |
|  |  | Age group x Sex | 0.001 | 0.001 | 1,19 | 1.054 | 0.318 |
|  |  | Age group x Level | 0.007 | 0.002 | 3,63 | 1.809 | 0.155 |
|  | Wave 2/3 | Age group | 0.012 | 0.012 | 1,19 | 1.154 | 0.296 |
|  |  | Sex | 0.022 | 0.022 | 1,19 | 2.015 | 0.172 |
|  |  | **Level** | **0.494** | **0.166** | **3,63** | **15.54** | **1.1e-7** |
|  |  | Age group x Sex | 0.018 | 0.018 | 1,19 | 1.664 | 0.213 |
|  |  | Age group x Level | 0.029 | 0.009 | 3,63 | 0.902 | 0.445 |
|  | Wave 4 | Age group | 0.000 | 1.0e-6 | 1,19 | 1.0e-4 | 0.992 |
|  |  | Sex | 0.025 | 0.025 | 1,19 | 1.926 | 0.181 |
|  |  | **Level** | **0.554** | **0.185** | **3,63** | **14.13** | **3.8e-7** |
|  |  | Age group x Sex | 0.004 | 0.004 | 1,19 | 0.274 | 0.607 |
|  |  | Age group x Level | 0.081 | 0.027 | 3,63 | 2.068 | 0.113 |
|  | Wave 5 | Age group | 0.015 | 0.015 | 1,19 | 0.312 | 0.583 |
|  |  | Sex | 0.013 | 0.012 | 1,19 | 0.254 | 0.619 |
|  |  | **Level** | **3.219** | **1.073** | **3,63** | **21.92** | **7.7e-10** |
|  |  | Age group x Sex | 6.0e-4 | 6.0e-4 | 1,19 | 0.012 | 0.913 |
|  |  | Age group x Level | 0.003 | 0.001 | 3,63 | 0.023 | 0.995 |
| ***24 kHz*** | | | | | | | |
| Wave 1 | | Age group | 1.4e-4 | 1.4e-4 | 1,20 | 0.146 | 0.706 |
|  |  | Sex | 1.4e-3 | 1.4e-3 | 1,20 | 1.482 | 0.238 |
|  |  | **Level** | **0.731** | **0.244** | **3,66** | **255.0** | **<2.0e-16** |
|  |  | Age group x Sex | 0.001 | 0.001 | 1,20 | 1.192 | 0.288 |
|  |  | Age group x Level | 0.007 | 0.002 | 3,66 | 2.392 | *0.076* |
| Wave 2/3 | | Age group | 5.5e-4 | 5.5e-4 | 1,20 | 0.157 | 0.696 |
|  |  | Sex | 3.8e-3 | 3.8e-3 | 1,20 | 1.071 | 0.313 |
|  |  | **Level** | **0.874** | **0.291** | **3,66** | **83.04** | **<2.0e-16** |
|  |  | Age group x Sex | 0.001 | 0.001 | 1,20 | 0.236 | 0.633 |
|  |  | Age group x Level | 0.008 | 0.003 | 3,66 | 0.768 | 0.516 |
| Wave 4 | | Age group | 0.006 | 0.006 | 1,20 | 0.376 | 0.547 |
|  |  | Sex | 0.055 | 0.055 | 1,20 | 3.337 | 0.087 |
|  |  | **Level** | **0.809** | **0.270** | **3,66** | **16.30** | **5.0e-8** |
|  |  | Age group x Sex | 0.002 | 0.002 | 1,20 | 0.147 | 0.706 |
|  |  | Age group x Level | 0.077 | 0.026 | 3,66 | 1.561 | 0.207 |
| Wave 5 | | Age group | 2.5e-3 | 2.5e-3 | 1,20 | 0.172 | 0.683 |
|  |  | Sex | 1.0e-3 | 1.0e-3 | 1,20 | 0.071 | 0.792 |
|  |  | **Level** | **1.604** | **0.534** | **3,66** | **37.27** | **3.3e-14** |
|  |  | Age group x Sex | 0.021 | 0.021 | 1,20 | 1.493 | 0.236 |
|  |  | Age group x Level | 0.025 | 0.008 | 3,66 | 0.591 | 0.623 |
| ***32 kHz*** | | | | | | | |
| Wave 1 | | Age group | 1.0e-4 | 9.9e-5 | 1,20 | 0.046 | 0.833 |
|  |  | Sex | 0.000 | 3.0e-6 | 1,20 | 0.001 | 0.971 |
|  |  | **Level** | **0.717** | **0.239** | **3,66** | **110.8** | **<2.0e-16** |
|  |  | Age group x Sex | 0.004 | 0.004 | 1,20 | 1.975 | 0.175 |
|  |  | Age group x Level | 0.008 | 0.003 | 3,66 | 1.339 | 0.269 |
| Wave 2/3 | | Age group | 2.0e-5 | 2.1e-5 | 1,20 | 0.004 | 0.953 |
|  |  | Sex | 1.1e-4 | 1.1e-4 | 1,20 | 0.018 | 0.895 |
|  |  | **Level** | **0.790** | **0.263** | **3,66** | **43.29** | **1.4e-15** |
|  |  | Age group x Sex | 0.005 | 0.005 | 1,20 | 0.843 | 0.370 |
|  |  | Age group x Level | 0.029 | 0.010 | 3,66 | 1.621 | 0.193 |
| Wave 4 | | Age group | 0.007 | 0.007 | 1,20 | 1.435 | 0.245 |
|  |  | Sex | 5.3e-4 | 5.3e-4 | 1,20 | 0.113 | 0.740 |
|  |  | **Level** | **1.010** | **0.337** | **3,66** | **71.51** | **<2.0e-16** |
|  |  | Age group x Sex | 2.7e-4 | 2.7e-4 | 1,20 | 0.057 | 0.813 |
|  |  | Age group x Level | 0.024 | 0.008 | 3,66 | 1.706 | 0.174 |
| Wave 5 | | Age group | 0.055 | 0.055 | 1,20 | 1.076 | 0.312 |
|  |  | Sex | 0.018 | 0.018 | 1,20 | 0.357 | 0.557 |
|  |  | **Level** | **1.610** | **0.537** | **3,66** | **10.54** | **9.4e-6** |
|  |  | Age group x Sex | 0.008 | 0.008 | 1,20 | 0.155 | 0.698 |
|  |  | Age group x Level | 0.219 | 0.073 | 3,66 | 1.433 | 0.241 |
|  | ***48 kHz*** | | | | | | |
|  | Wave 1 | Age group | 0.004 | 0.004 | 1,20 | 0.431 | 0.519 |
|  |  | Sex | 0.002 | 0.002 | 1,20 | 0.219 | 0.645 |
|  |  | **Level** | **0.629** | **0.210** | **3,64** | **25.30** | **6.4e-11** |
|  |  | Age group x Sex | 0.020 | 0.020 | 1,20 | 2.396 | 0.137 |
|  |  | Age group x Level | 0.024 | 0.008 | 3,64 | 0.973 | 0.411 |
|  | Wave 2/3 | Age group | 8.8e-4 | 8.8e-4 | 1,20 | 0.037 | 0.849 |
|  |  | Sex | 2.2e-3 | 2.2e-3 | 1,20 | 0.094 | 0.762 |
|  |  | **Level** | **1.523** | **0.508** | **3,64** | **21.33** | **1.1e-9** |
|  |  | Age group x Sex | 0.036 | 0.036 | 1,20 | 1.519 | 0.232 |
|  |  | Age group x Level | 0.031 | 0.010 | 3,64 | 0.429 | 0.732 |
|  | Wave 4 | Age group | 0.001 | 0.001 | 1,20 | 0.037 | 0.850 |
|  |  | Sex | 2.2e-4 | 2.1e-4 | 1,20 | 0.007 | 0.937 |
|  |  | **Level** | **0.912** | **0.304** | **3,64** | **9.355** | **3.3e-5** |
|  |  | Age group x Sex | 0.023 | 0.023 | 1,20 | 0.714 | 0.408 |
|  |  | Age group x Level | 0.036 | 0.012 | 3,64 | 0.370 | 0.775 |
|  | Wave 5 | Age group | 0.000 | 0.000 | 1,20 | 0.000 | 0.995 |
|  |  | Sex | 3.0e-5 | 3.0e-5 | 1,20 | 0.001 | 0.978 |
|  |  | **Level** | **1.155** | **0.385** | **3,64** | **8.996** | **4.7e-5** |
|  |  | Age group x Sex | 0.131 | 0.131 | 1,20 | 3.052 | 0.096 |
|  |  | Age group x Level | 0.058 | 0.019 | 3,64 | 0.453 | 0.716 |

Results of linear mixed-effects modeling testing the significance of fixed effects (age group, level, and sex and the interactions of age group and sex and age group and level)) on the measured latencies of ABRs evoked by 20-48 kHz tones. For all tests, subject ID was included as a random effect to account for individual variation.

**Table S6. Correlation of ABR wave 1 amplitudes and latencies with chronological age**

| **Stimulus** | **Level** | **Pearson correlation coefficient (r)** | **t-value** | **df** | **p-value** |
| --- | --- | --- | --- | --- | --- |
| *Amplitude* |  |  |  |  |  |
| Click | 60 dB | 0.305 | 1.469 | 21 | 0.157 |
|  | 70 dB | 0.041 | 0.186 | 21 | 0.854 |
|  | 80 dB | 0.029 | 0.134 | 21 | 0.894 |
|  | 90 dB | -0.052 | -0.243 | 22 | 0.810 |
| 20 kHz | 60 dB | 0.181 | 0.845 | 21 | 0.407 |
|  | 70 dB | 0.370 | 1.826 | 21 | 0.082 |
|  | 80 dB | 0.340 | 2.247 | 21 | 0.096 |
|  | 90 dB | 0.299 | 1.437 | 21 | 0.165 |
| 24 kHz | 60 dB | -0.047 | -0.221 | 22 | 0.827 |
|  | 70 dB | 0.030 | 0.140 | 22 | 0.890 |
|  | 80 dB | 0.003 | 0.016 | 22 | 0.988 |
|  | 90 dB | 0.012 | 0.055 | 22 | 0.956 |
| 32 kHz | 60 dB | 0.024 | 0.114 | 22 | 0.911 |
|  | 70 dB | 0.067 | 0.314 | 22 | 0.757 |
|  | 80 dB | -0.088 | -0.415 | 22 | 0.682 |
|  | 90 dB | 0.082 | 0.386 | 22 | 0.704 |
| 48 kHz | 60 dB | -0.086 | -0.385 | 20 | 0.705 |
|  | 70 dB | 0.110 | 0.520 | 22 | 0.609 |
|  | 80 dB | 0.078 | 0.366 | 22 | 0.718 |
|  | 90 dB | 0.067 | 0.316 | 22 | 0.755 |
| *Latency* |  |  |  |  |  |
| Click | 60 dB | 0.047 | 0.217 | 21 | 0.831 |
|  | 70 dB | -0.132 | -0.608 | 21 | 0.549 |
|  | 80 dB | -0.103 | -0.475 | 21 | 0.640 |
|  | 90 dB | -0.268 | -1.306 | 22 | 0.205 |
| 20 kHz | 60 dB | 0.108 | 0.497 | 21 | 0.624 |
|  | 70 dB | 0.115 | 0.532 | 21 | 0.600 |
|  | 80 dB | 0.117 | 0.538 | 21 | 0.596 |
|  | 90 dB | -0.112 | -0.517 | 21 | 0.611 |
| 24 kHz | 60 dB | -0.126 | -0.594 | 22 | 0.559 |
|  | 70 dB | -0.127 | -0.601 | 22 | 0.554 |
|  | 80 dB | -0.054 | -0.255 | 22 | 0.801 |
|  | 90 dB | -0.049 | -0.228 | 22 | 0.822 |
| 32 kHz | 60 dB | -0.124 | -0.586 | 22 | 0.564 |
|  | 70 dB | -0.059 | -0.278 | 22 | 0.784 |
|  | 80 dB | 0.011 | 0.053 | 22 | 0.959 |
|  | 90 dB | 0.145 | 0.687 | 22 | 0.499 |
| 48 kHz | 60 dB | -0.147 | -0.662 | 20 | 0.515 |
|  | 70 dB | -0.296 | -1.456 | 22 | 0.160 |
|  | 80 dB | -0.112 | -0.530 | 22 | 0.602 |
|  | 90 dB | -0.147 | -0.699 | 22 | 0.492 |
